# High-content fluorescence bioassay to investigate pore formation, ion channel modulation and cell membrane lysis induced by venoms

**DOI:** 10.1101/2023.10.25.563942

**Authors:** Simon Kramer, Charan Kotapati, Yuanzhao Cao, Bryan G Fry, Nathan J Palpant, Glenn F King, Fernanda C Cardoso

## Abstract

Venoms comprise highly evolved bioactive molecules modulating ion channels, receptors, coagulation factors, and the cellular membrane. This array of targets and bioactivity requires high-content bioassays to aid the development of novel envenomation treatments and biotechnological and pharmacological agents. To address this gap in venom’s research, we developed a fluorescence-based high-throughput and high-content cellular assay to simultaneously identify common cellular activities produced by venoms: membrane lysis, pore-formation, and ion channel modulation. By combining intracellular calcium with extracellular nucleic acid measurements, we distinguished these venom mechanisms using one cellular assay. We applied our high-content bioassay in three cell types exposed to venom components representing lytic, ion pore-forming or ion channel modulator toxins. Beyond the distinct profiles produced by these three types of action mechanisms, we found that the pore-forming latrotoxin α-Lt1a prefers human neuroblastoma to kidney cells and cardiomyocytes, while the lytic bee peptide melittin is not selective. Furthermore, evaluation of snake venoms showed that Elapid species induced rapid membrane lysis, while Viper species showed variable to no activity on neuroblastoma cells. These findings demonstrate that our high-content bioassay distinguish clades and interspecific traits and capture the clinical observations at venom level and is capable of differentiate ion pore-forming from membrane lysis and ion channel modulation. We hope our research will accelerate the understanding of venom biology and the diversity of toxins inducing cytotoxic, cardiotoxic and neurotoxic effects and assist in identifying venom components whose properties could benefit humankind.

## Introduction

Venoms are unique cocktails of bioactive molecules with exquisite properties to modulate human physiology. In nature, venoms can induce severe harm to the envenomated through neurotoxic, hemotoxic, and cytotoxic effects^1^. Fortunately, research in venoms has found that toxins in isolation can elicit unique pharmacological properties with applications in therapeutics and biopesticides^2,3^. The mechanisms of action and pharmacological properties of many venom components have been elucidated, unrevealing a complex multifunctional cocktail acting synergistically to immobilize prey and deter predators^1,4,5^. Primitive mechanisms in venom biology include cell membrane lysis and cytolysis, as observed in bee venom^6^, while sophisticated envenomation mechanisms induce ion pore-formation and ion channel/receptor modulation, as in spider venoms^4,7^. The understanding of these mechanisms is critical for developing more efficacious envenomation treatments^8^, and identifying bioactive entities with commercial potential.

The first drug derived from an animal venom is captopril, an inhibitor of the angiotensin-converting enzyme used in hypertension and congestive heart failure^9,10^. It is derived from proline-rich oligopeptides from the venom of the Brazilian snake *Bothrops jararaca*^11,12^. This translational science milestone in the 70s revealed the exceptional potential of toxins as a novel source of bioactive molecules for drug development. Since then, at least 11 novel drugs derived from venoms became available for various conditions, including hypertension, chronic pain and diabetes type II^3^.

Progress in venom-based drug discovery depends on robust and reliable high-throughput cellular assays to identify venom components’ function in the early stages of the discovery process. Amongst popular assay read-outs is fluorescence-based intracellular calcium measurement that are low-cost, readily accessible, and produce robust data. Calcium ions play a central role in various biological functions and cellular processes which facilitates the identification of many biological processes but that also challenges interpretation of bioactive-rich mixtures like venoms. A major caveat is that venoms usually contain protein and peptides that lysis the cell membrane, making the relevance of calcium responses questionable as cytolytic effects trigger a rapid rise of intracellular calcium levels, which can be mistaken with other cellular events such as receptor and ion channel modulation^13^.

In this work, we developed a duplex fluorescence assay to monitor ion channel activity and cytolytic effects via a combination of intracellular calcium nd extracellular DNA measurements. We found that intracellular calcium dye combined with the DNA marker propidium iodide allowed simultaneous measurement of intracellular calcium responses and DNA exposure in real-time recording and produced profiles that differentiated membrane lysis from ion pore-formation and ion channel modulation. This new high-content bioassay was applied to studies of snake venom biology, showing its usefulness in phylogenetic and venom bioactivity investigations. This article is a guide for studies of venom-based drug discovery and venom biology and optimization of high-throughput into high-content bioassays.

## Materials and Methods

### Reagents

Reagents were from Gibco (MA, USA) and SIGMA (MO, USA). Calcium 4 dye was from Molecular devices (CA, USA). Assay buffer (physiology saline solution - PSS) contained (in millimolar) 140 NaCl, 11.5 glucose, 5.9 KCl, 1.4 MgCl_2_, 1.2 NaH_2_PO_4_, 5 NaHCO_3_, 1.8 CaCl_2_, and 10 HEPES (pH 7.4). PSS buffer with 0.1% BSA was used in all reagents for the assays. Native α-latrotoxin-Lt1a was purchased from Alomone (Jerusalem, ISR), synthetic melittin and OD1 were purchased from Smartox Biotechnology (Saint Egrève, FRA).

### Cell lines

SHSY5Y neuroblastoma cells were cultured in RPMI supplemented with 15% FBS, 2 mM Glutamine and 100 units.ml^-1^ Penicillin 100 μg.ml^-1^ Streptomycin. HEK293 cells were cultured in DMEM supplemented with 10% FBS, and 100 units.ml^-1^ Penicillin 100 μg.ml^-1^ Streptomycin. Cells were incubated at 37°C in a humidified 5% CO_2_ incubator and subcultured every 2-3 days in a 1:5 ratio using 0.05% Trypsin/EDTA and D-PBS after 70-80% confluency. hiPSC cardiomyocytes were prepared as previously described by us^14^. Briefly, on day -1 of differentiation, hiPSC were dissociated using 0.5% EDTA, plated onto Vitronectin XF-coated flasks (Nunc, USA) at a density of 1.12 × 10^5^ cells/cm^2^, and cultured overnight in mTeSR Plus medium supplemented with 10 µM Y-27362 ROCK inhibitor (Stem Cell Technologies, Canada). Differentiation was induced on day 0 by first washing cells with PBS when the monolayer reached approximately 80% confluence, then changing the culture medium to RPMI 1640 Medium (ThermoFisher, MA, USA) containing 3 μM CHIR99021 (Stem Cell Technologies, Canada), 500 μg.mL^-1^ bovine serum albumin (BSA, Sigma Aldrich), and 213 μg/mL ascorbic acid (AA, Sigma Aldrich). After 3 days of culture, the medium was replaced with RPMI 1640 containing 5 μM XAV-939 (Stem Cell Technologies, Canada), 500 μg.mL^-1^ BSA, and 213 μg.mL^-1^ AA. On day 5, the medium was exchanged to RPMI 1640 containing 500 μg.mL^-1^ BSA, and 213 μg.mL^-1^ AA without additional supplements. From day 7 and onward, the cultures were fed every other day with RPMI 1640 containing 2% B27 supplement with insulin (Life Technologies, Australia).

### Venoms

Crude venoms from snakes were kindly provided by collaborators or purchased from SIGMA-Aldrich. Venom from *Pseudechis porphyriacus* and *Pseudechis australis* were provided by Bryan Fry at the Venom Evolution laboratory in the University of Queensland, AU. Venoms from *Bothrops jararaca* and *Crotalus horridus* were provided by Camila Ferraz at the State University of Londrina, BR. The venom of *Echis carinatus* was purchased from Sigma-Aldrich. Venoms were stored in -80 °C and weighed and reconstituted in PSS buffer just before bioactivity analysis. After reconstitution, crude venoms were further quantified using Nanodrop 2000c (Thermofisher, MA, USA).

### Fluorescence-imaging duplex assay

Hight throughput cellular assays were performed using the FLIPR Penta System (Molecular Devices). Cells were seeded in 384 wells black wall and flat clear back bottom microplate (Corning, NY, USA) at 40,000 cells.well^-1^ for SHSY5Y and 10,000 cells.well^-1^ for HEK293, and incubated at 37**°**C in a humidified 5% CO_2_ incubator for 48hrs before the assays. HiPSC cardiomyocytes were replated on day 15 of differentiation. Cells were dissociated using 0.5% trypsin, stopped with Stop Buffer (1:1 FBS in RPMI 1640), filtered with a 100 μm strainer, replated at a density of 10,000 cells/well in Vitronectin XF-coated black-walled, clear-bottom Cellbind 384-well microplates (Corning, NY, USA) with replating medium (RPMI 1640, 2% B27 supplement plus insulin, 5% FBS, and 10 µM Y-27362) and cultured overnight. FBS and ROCK inhibitor-containing medium was replaced the following day (day 0) with standard medium (RPMI 1640 plus B27/insulin). From day 0 to day 7 post-replate, cells were refreshed with standard medium every other day before the assay date.

On the assay day, cells were loaded with 20 μl per well of propidium iodide in Calcium 4 dye reconstituted in PSS and incubated for 30 min at 37**°**C in a humidified 5% CO_2_ incubator. Fluorescence responses for read mode 1 (Propidium iodide) were recorded at 470-495 nm excitation and 565-625 nm emission, while fluorescence responses for read mode 2 (Calcium 4 dye) were recorded at 470-495 nm excitation and 515-575 nm emission. The baseline of 5 (SHSY5Y and HEK293) to 100 (cardiomyocytes) readings before dispensing reagents to the cells was used to set the baseline, followed by readings after addition of toxins or crude venoms to equal a total reading time of up to 100 minutes, and when applicable readings after KCl 90 mM plus CaCl_2_ 5 mM exposure to equal a total time of 5 mins.

### Data analysis

Fluorescence intensity kinetic reduction was measured as both Max-Min and Area Under the Curve (AUC) from the highest response normalised against positive control or highest response, and negative controls (buffer only). Z-scores were calculated using ScreenWorks® Software (Molecular Devices). All graphs were generated using GraphPad Prism 9.0. Curve fitting was performed using nonlinear regression with log-inhibitor versus normalized response and variable Hill slope for dose-responses and EC_50_ determination.

## Results

### Assay development

We evaluated the intracellular calcium responses and the extracellular DNA exposure due to membrane lysis using a duplex fluorescence assay containing 30 μM or 50 μM of propidium iodide (PI) mixed in Calcium 4 dye (Ca4). Triton X-100 (1 %) added to the cells induced a transient increase in [Ca^2+^]_i_ and increased and sustained DNA exposure response (Figure 1A); the PSS buffer used as negative control produced no modification in intracellular calcium or DNA exposure. Interestingly, 30 μM or 50 μM PI in the absence of Ca4 did not alter the fluorescence intensity for DNA labelling, but in combination with Ca4 dye the maximum fluorescence intensity for DNA labelling increased by 2 to 3-fold (Figure 1B). The membrane lysis induced by Triton produced a transient [Ca^2+^]_i_ rise followed by complete depletion of Ca^2+^ signals over time due to the presence of an extracellular quencher for the Ca^2+^ dye (masking technology, Molecular Devices), while DNA exposure responses was sustained (Figure 1C).

**Figure 1.**
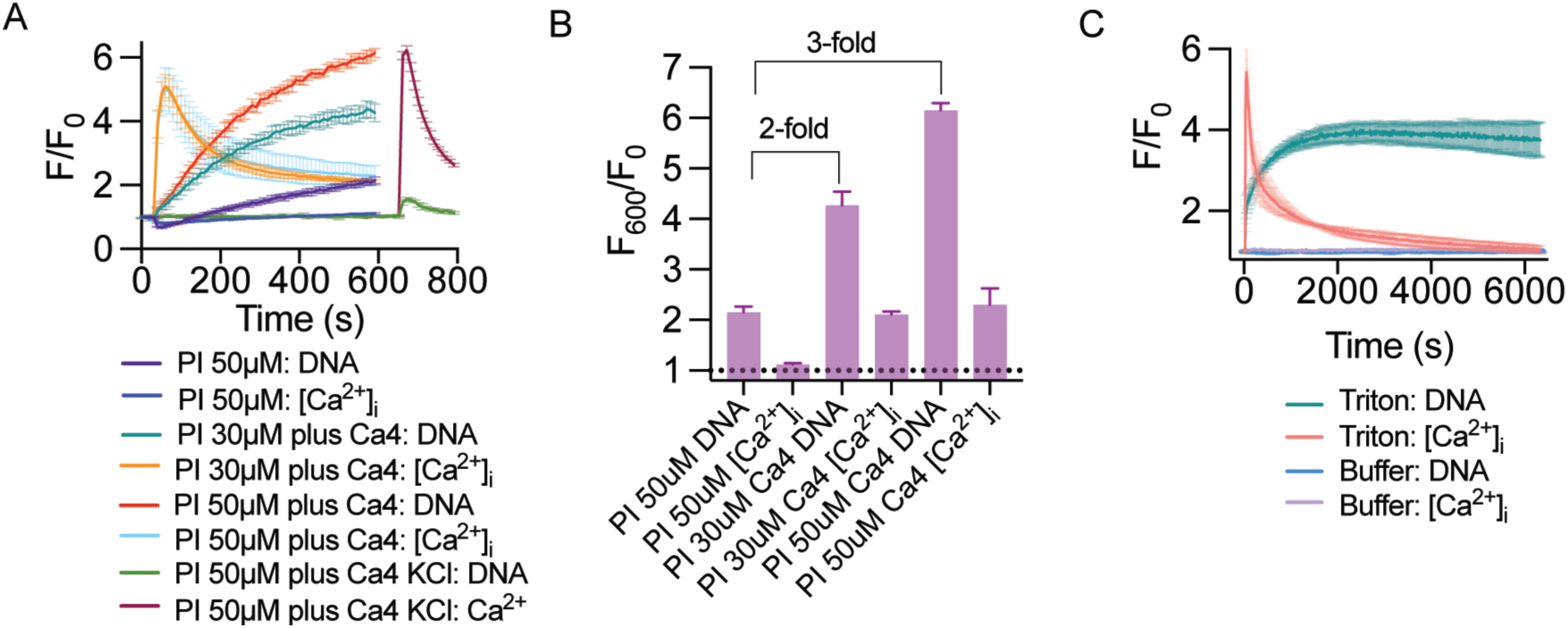
Duplex fluorescence assay in neuroblastoma SHSY5Y cells in the presence of Triton X-100. (A) Fluorescence traces of SHSY5Y cells in the presence of 1 % Triton X-100 and 30 μM or 50 μM of propidium iodide (PI, DNA) and calcium 4 dye (Ca4, [Ca^2+^]_i_). KCl 90 mM was added after 600 s to evoke Ca_V_ responses in the presence of the dye duplex and absence of Triton. (B) Absolute fluorescence values at 600s represented by bar graphs showing a 2 to 3-fold increase in PI plus DNA signals in the presence of Ca4, with [Ca^2+^]_i_ not altered in the dye duplex (PI plus Ca4). (C) Representative fluorescence traces of 50 μM PI and Ca4 dye signals in the presence of 1 % Triton X-100 in SHSY5Y cells. Traces showed the DNA response is sustained and the [Ca^2+^]_i_ is transient, consistent with the profile of cell lysis. Data are represented by mean ± SEM from triplicates.

The concentration of 50 μM Propidium iodide plus Calcium 4 dye exhibited the maximum signals for DNA detection (Figure 1A). Therefore, we used 50 μM Propidium iodide mixed in Calcium 4 dye for all subsequent evaluations in this study. The concentration of 100 μM Propidium iodide plus Calcium 4 dye exhibited similar intensity signal for DNA labelling to 50 μM (data not shown). The Z factor for this assay was calculated via propidium iodide fluorescence and showed a value of 0.81 from n=12 independent wells in the presence of Triton X-100 1%. The same wells measured via Calcium 4 dye showed a Z factor of 0.63.

The Calcium 4 dye employs a masking technology that significantly reduces background fluorescence originating from residual extracellular calcium indicator, media and other components (Molecular Devices). Despite this, background response was present when voltage-gated calcium channel (Ca_V_) responses were evoked by KCl (Figure 1A). The Ca_V_ response produced a high and transient rise in [Ca^2+^]_i_ and a weak and transient PI response. An explanation for this unexpected PI response is an overlap of 10 nm within the FLIPR filters wavelengths of emission for the propidium iodide read mode (565-625 nm) and the calcium 4 dye (515-575 nm). This produces a small leak of calcium 4 dye fluorescence into the PI channel. Filter with defined ranges for this fluorescence duplex assay are recommended to eliminate this artefact leak response.

### Membrane lysis induced by melittin

Melittin, a bee venom peptide known by its properties that lysis the cell membrane^15^, induced a dose-dependent transient [Ca^2+^]_i_ increase and sustained DNA response characteristic of formation of large pores permeable to DNA and cytolysis in SHSH5Y cells (Figure 2A), and which resembled the positive control Triton-induced cytolysis. Dose-response curves produced EC_50_ values of 2.2 and 0.57 μM for DNA exposure and [Ca^2+^]_i_ increase, respectively, in SHSY5Y cells (Figure 2B, Table 1), while in HEK293 cells the melittin-induced membrane lysis and cytolysis produced EC_50_ values of 3.8 and 5.3 μM for DNA exposure and [Ca^2+^]_i_ increase, respectively (Figure 2C, Table 1). These results suggested melittin induced a more potent [Ca^2+^]_i_ increase in neuroblastoma, potentially via ion channel or receptor modulation (13-fold more potent compared to DNA release, Figure 2G), which is rapidly overtaken by the membrane lysis process as showed by the sustained DNA signal (Figure 2A, B, and G). The same is not observed in HEK293 cells that showed a similar EC_50_ for DNA release and [Ca^2+^]_i_ increase (Figure 2C).

**Figure 2.**
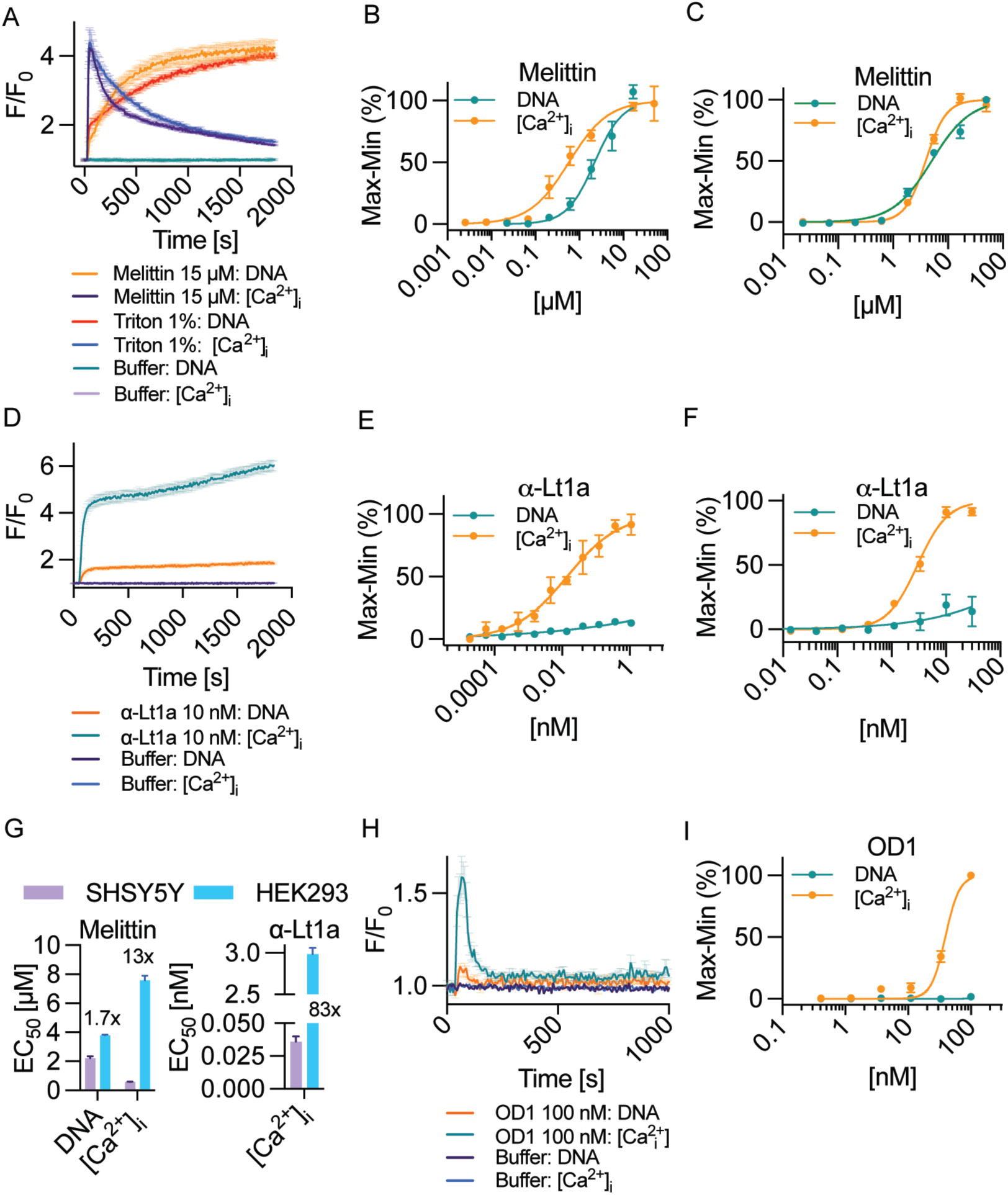
Duplex fluorescence assay using human neuroblastoma SHSY5Y and HEK293 cells in the presence of synthetic or purified venom components representing membrane lysis, pore-formation, or ion channel modulation. (A-C) Representative fluorescence traces of DNA exposure and [Ca^2+^]_i_ measurements in the presence of 15 μM melittin or 1 % Triton X-100 (A) demonstrating membrane lysis and cytolysis, and calculated dose responses for the SHSY5Y (B) and HEK293 cells (C). (D) Representative fluorescence traces of DNA exposure (PI) and intracellular calcium (Ca4) measurements in the presence of 10 nM α-Lt1a, and calculated dose responses for the SHSY5Y (E) and HEK293 cells (F). (G) Comparison of the EC_50_ values calculated for melittin and α-Lt1a for the SHSY5Y and HEK293 cells. While melittin showed potent induction of [Ca^2+^]_i_ increase at up 13x for SHSY5Y, the α-Lt1a showed a striking increase of 83x potency for SHSY5Y compared to HEK293. (H) Representative fluorescence traces of DNA exposure and [Ca^2+^]_i_ measurements in the presence of 100 nM OD1 and calculated dose responses for the ion channel modulatory effects on SHSY5Y cells (I). EC_50_ values are described in Table 1. Data are represented as mean ± SEM of triplicates (n=3 independent wells).

**Table 1.** Calculated EC_50_ values for the dose response curves in the presence of isolated venom toxins evaluated on human neuroblastoma SHSY5Y or HEK293 cells using the duplex fluorescence assay. EC_50_ values are represented in various units from pM to μM. Data are represented as mean ± SEM of triplicates (n=3 independent wells).

| Venom peptide | SHSY5Y |  | HEK293 |  |
| --- | --- | --- | --- | --- |
| | $\text{EC}_{50}$ DNA | $\text{EC}_{50}$ $[\text{Ca}^{2+}]_i$ | $\text{EC}_{50}$ DNA | $\text{EC}_{50}$ $[\text{Ca}^{2+}]_i$ |
| α- Lt1a | n.a. | $36 \pm 0.4$ pM | n.a. | $3 \pm 0.07$ nM |
| Melittin | $2.2 \pm 0.13$ $\mu\text{M}$ | $576 \pm 38$ nM | $3.8 \pm 0.06$ $\mu\text{M}$ | $5.3 \pm 0.17$ $\mu\text{M}$ |
| OD1 | n.a. | $40 \pm 1$ nM | n.a. | n.a. |

### Ion pore formation and ion channel modulation induced by venom toxins

The spider toxin α-Lt1a is the major neurotoxin responsible for the human envenomation syndrome latrodectism^16^ and a well-characterized Ca^2+^ ion permeable pore-forming toxin^17^. In this study, α-Lt1a induced a dose-dependent sustained [Ca^2+^]_i_ increase and no DNA release responses consistent with the Ca^2+^ ion pore-formation in SHSY5Y cell membranes (Figure 2D). Dose-response curves produced EC_50_ values of 36 pM for [Ca^2+^]_i_ increase in SHSY5Y (Figure 2E), while in HEK293 cells the α-Lt1a induced ion pore formation with EC_50_ values of 3 nM for [Ca^2+^]_i_ increase (Figure 2F). These results indicated α-Lt1a has higher affinity for the neuroblastoma cell line, with at least 83x more potency to modulate SHSY5Y compared to the HEK293 cells (Figure 2G). The scorpion toxin OD1, known by its properties to activate voltage-gated sodium channels (Na_V_)^18^, induced a dose-dependent transient [Ca^2+^]_i_ increase and no DNA responses consistent with the activation of voltage-gated ion channels in SHSH5Y cells (Figure 2H). Dose-response curves produced EC_50_ values of 40 nM [Ca^2+^]_i_ increase for the SHSY5Y cells (Figure 2B). This toxin was not tested on HEK293 cells due to the absence of endogenous expression of relevant voltage-gated ion channels in this cell line.

### Human iPSC cardiomyocytes response to venom toxins

Human iPSC cardiomyocytes produced similar responses to Triton, melittin and α-Lt1a compared to SHSY5Y and HEK293 cells, while OD1 accelerated the [Ca^2+^]_i_ spiking characteristic of the muscle contraction in this cell type^19^ (Figure 3).

**Figure 3.**
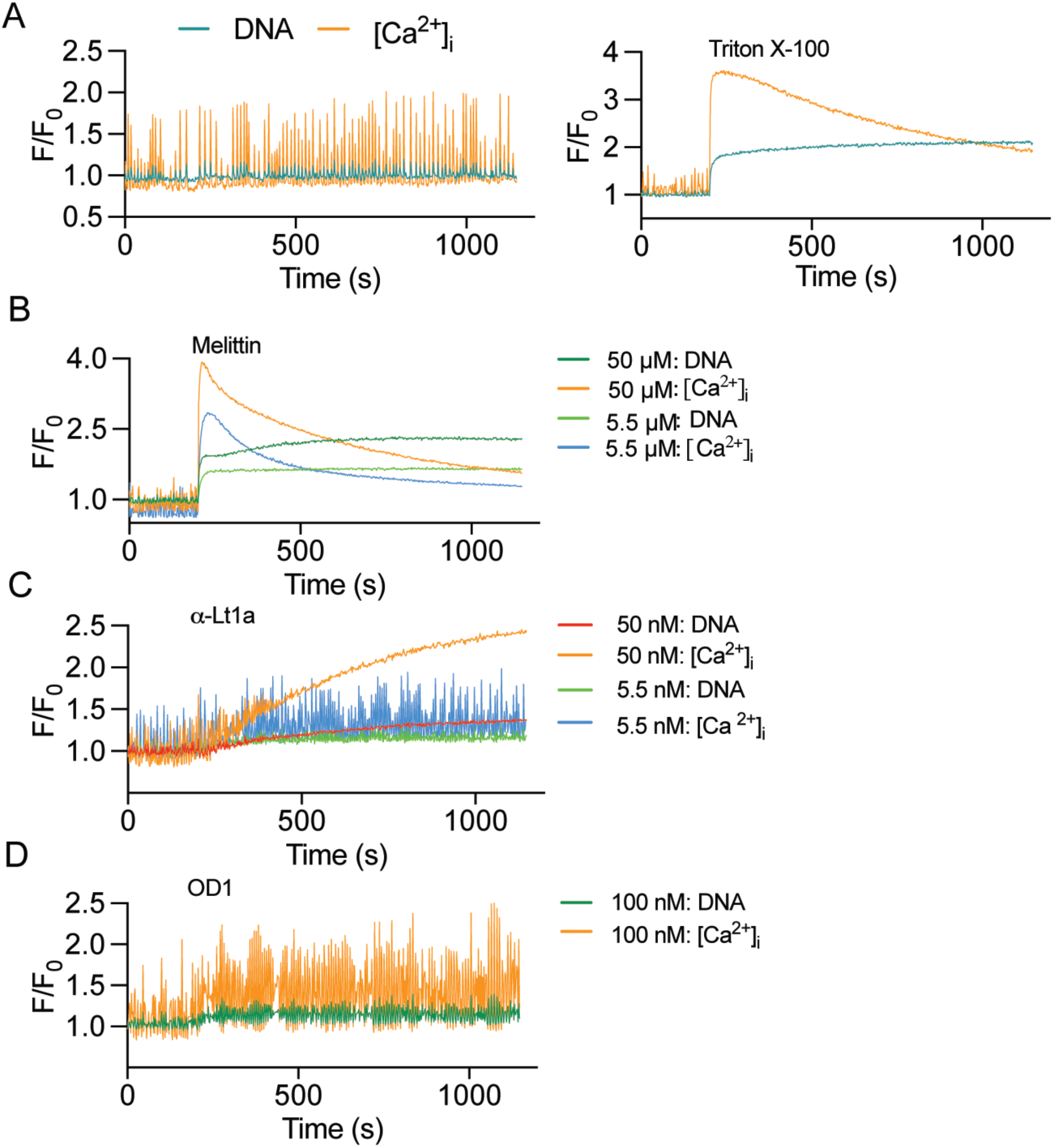
Duplex fluorescence assay using hiPSC cardiomyocytes in the presence of isolated venom toxins. (A) Representative fluorescence traces of DNA exposure and [Ca^2+^]_i_ measurements in cardiomyocytes in the presence of buffer only (left graph) showing the characteristic [Ca^2+^]_i_ increase spiking regenerate by the cardiac muscle contraction; or in the presence of 1% Triton (right graph) showing the characteristic DNA exposure and transient [Ca^2+^]_i_ increase typical of membrane lysis and cytolysis. (B) Melittin at 50 or 5.5 μM induced membrane lysis and cytolysis with DNA exposure and transient [Ca^2+^]_i_ increase. We observed a mixed response of ion pore formation and DNA release at 1.8 μM melittin and non-altered cellular responses at 617 nM melittin (data not shown). (C) Representative fluorescence traces of DNA exposure and [Ca^2+^]_i_ measurements in the presence of 50 or 5.5 nM α-Lt1a showing a typical response of a ion pore formation with sustained [Ca^2+^]_i_ signals and absence of DNA exposure. (D) Representative fluorescence traces of DNA exposure and [Ca^2+^]_i_ measurements in the presence of 100 nM OD1 showing an increase in the frequency of [Ca^2+^]_i_ spiking which is consistent with the Na_V_ channel modulatory effects of OD1. Data are represented by representative traces of each condition evaluated in triplicates (n=3 independent wells).

In the presence of Triton 1%, the cardiomyocytes produced a transient [Ca^2+^]_i_ increase and sustained DNA responses characteristic of membrane lysis and cytolysis (Figure 1A, right graph). Melittin had a similar response to Triton and produced sustained DNA release at various concentrations tested from 50 to 5.5 μM, indicating the formation of large pores permeable to DNA (Figure 3B). Interestingly, we observed a mixed response of ion pore formation and DNA release at 1.8 μM melittin and non-altered cellular responses at 617 nM (data not shown). The α-Lt1a at 50 nM induced a sustained [Ca^2+^]_i_ increase and no DNA release responses characteristic of Ca^2+^ ion pore-formation in cardiomyocytes (Figure 3C). A dose-response curve was not calculated with this cell line. At lower concentration of 5.5 nM, the α-Lt1a increased the [Ca^2+^]_i_ spiking frequency in absence of sustained [Ca^2+^]_i_ increase. Similarly, the scorpion peptide OD1 at 100 nM produced an increase in the [Ca^2+^]_i_ spiking frequency and absence of sustained [Ca^2+^]_i_ increase (Figure 3D). This response is consistent with the OD1 ability to activate Na_V_ channels, including the subtype Na_V_1.5 expressed in the heart tissue.

### Cytotoxicity induced by Elapid snake venoms

The cytotoxicity (also refereed as membrane lysis and cytolysis in this work) induced by crude venoms from Australian Elapid snakes *P. porphyriacus* (Red-bellied black snake) and *P. australis* (king brown snake) were evaluated using the duplex fluorescence assay with the SHSY5Y cells and described in Figure 4 and Table 2.

**Figure 4.**
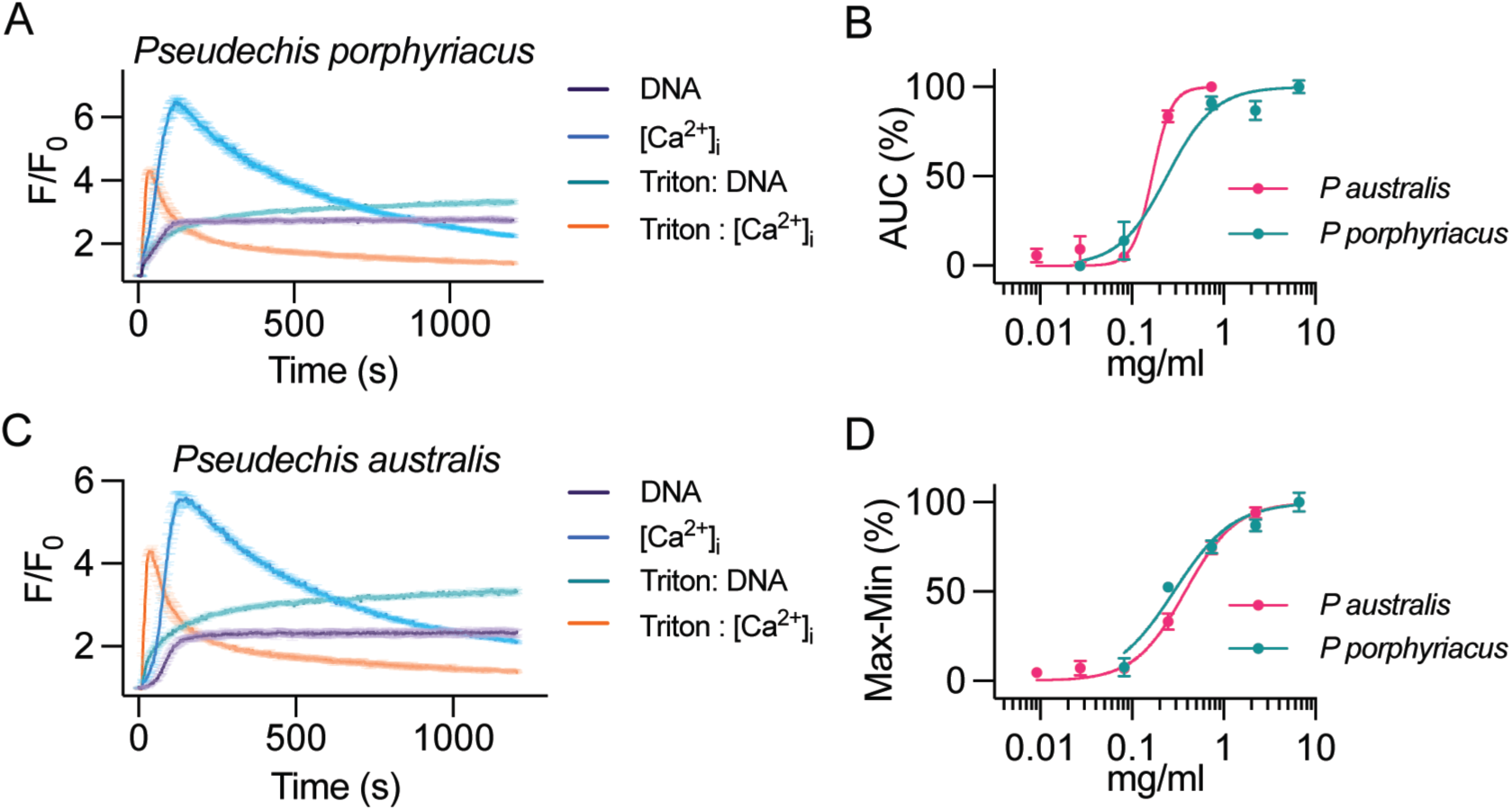
Snake Elapid venoms evaluation in neuroblastoma SHSY5Y cells using the duplex assay. (A and C) Representative fluorescence traces of DNA exposure and [Ca^2+^]_i_ responses in SHSY5Y cells in the presence of 2.2 mg/ml *P. porphyriacus* or *P. australis* venoms. These venoms induced typical responses membrane lysis by producing sustained DNA exposure and transient [Ca^2+^]_i_ increase. Triton 1% and buffer were used as controls. (B and D) Dose response curves of *P. porphyriacus* or *P. australis* snake crude venoms measured using propidium iodide (B, DNA) or calcium 4 dye (D, [Ca^2+^]_i_). Data are represented as mean ± SEM in a non-linear regression model (variable slope). The calculated EC_50_ values are described in Table 2.

**Table 2.** Calculated EC_50_ values for the cytotoxicity dose responses of crude venoms evaluated in human neuroblastoma using the duplex assay. Data are represented as mean ± SEM of one experiment in triplicate (n=3 independent wells).

| Snake species | EC<sub>50</sub> DNA<br>(µg/ml) | EC<sub>50</sub> [Ca<sup>2+</sup>]<sub>i</sub><br>(µg/ml) |
| --- | --- | --- |
| *P. porphyriacus* | 242 ± 25 | 286 ± 15 |
| *P. australis* | 167 ± 0.007 | 380 ± 0.011 |
| *E. carinatus* | 56410 ± 8796 | 647 ± 37 |
| *B. jararaca* | 2423 ± 80 | 1165 ± 70 |
| *C. horridus* | n.a. | n.a. |

In cytotoxin-rich Elapid venoms, DNA exposure measurements for *P. porphyriacus* and *P. australis* produced high and sustained responses to propidium iodide (Figure 4A and C) and transient [Ca^2+^]_i_ increase, which is characteristic of membrane lysis and cytolysis. The dose responses and the calculated EC_50_ values for the cytotoxicity induced by these venoms were for *P. porphyriacus* 242 and 286 μg/ml for DNA and [Ca^2+^]_i_ measurements, respectively, and for *P. australis* 167 and 380 μg/ml for DNA and [Ca^2+^]_i_ measurements, respectively (Figures 4B and D, Table 2). Consistent with the epidemiological studies showing the king brown snake venom is the most necrotic compared to the red-bellied black snake^20^, our results showed brown snake with the most potent venom (1.4-fold increased) to induce membrane lysis.

### Cytotoxicity induced by Viper snake venoms

The cytotoxic effects induced by crude venoms from Vipiride snakes *B. jararaca*, *E. carinatus* and *C. horridus* were evaluated on SHSY5Y cells and described in Figure 5. In contrast to Elapid snake venoms, Viperidae venoms are rich in phospholipases and proteases that induce cytotoxicity through a slower mechanism. The *B. jararaca* venom induced transient [Ca^2+^]_i_ increase and weak propidium iodide signal (Figure 5A), while *E. carinatus* produced a similar but weaker response to propidium iodide (Figure 5C). Interestingly, *C. horridus* venom did not induce propidium iodide or calcium 4 dye responses (Figure 5E).

**Figure 5.**
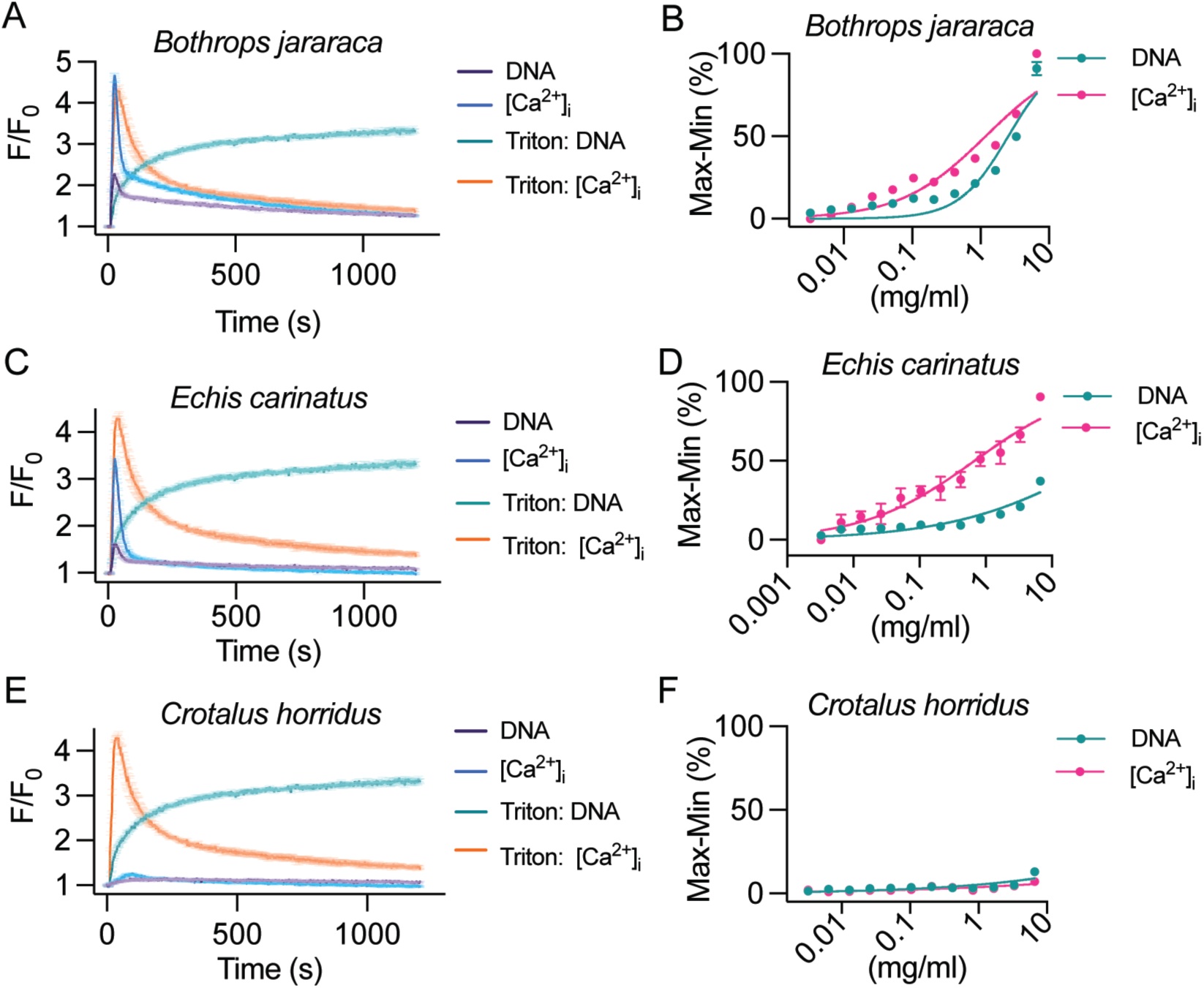
Snake Viper venoms cytotoxicity in neuroblastoma SHSY5Y cells using the duplex assay. (A, C and E) Representative fluorescence traces of DNA exposure and [Ca^2+^]_i_ responses in SHSY5Y cells in the presence of 3.3 mg/ml *B. jararaca, E. carinatus or C horridus* venoms. The venoms from B jararaca and E carinatus induced typical responses ion channels or receptors modulation by producing transient [Ca^2+^]_i_ increase in the absence of sustained DNA exposure. The venom of *C. horridus* did not alter the SHSY5Y cellular response. Triton 1% and buffer were used as controls. (B, D and F) Dose response curves of *B. jararaca, E. carinatus or C horridus* snake crude venoms measured using propidium iodide (DNA) or calcium 4 dye ([Ca^2+^]_i_). Data are represented as mean ± SEM in a non-linear regression model (variable slope). The calculated EC_50_ values are described in Table 2.

The dose responses and the EC_50_ for the cytotoxicity induced by these venoms are described in Figures 5B, D and F and in Table 2, with *E. carinatus* inducing the most potent [Ca^2+^]_i_ modulation within the vipers studies in this work and complete absence of DNA exposure, highlighting this snake venom for further characterization of potential neuroactive components. The ranking of cytolytic activity for the snake venoms studied in this work is represented in Figure 6, with *P. australis* being the most cytolytic and *C. horridus* the least cytolytic venom.

**Figure 6.**
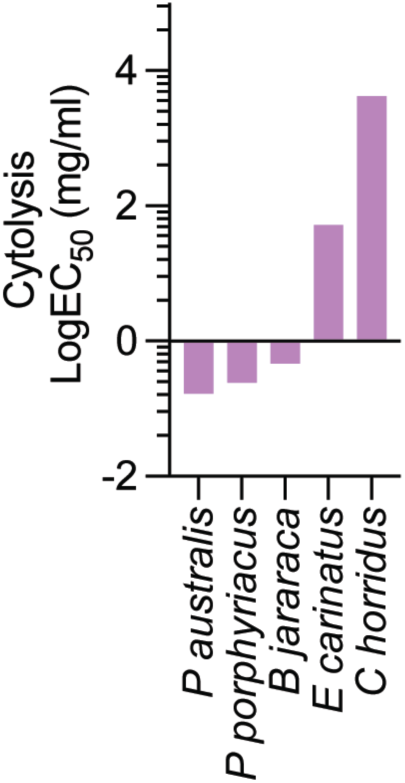
Comparison of the LogEC_50_ values calculated for the DNA exposure response induce by snake venoms. The ranking of the potency of membrane lysis by snake venoms confirmed the elapid venoms from *P. australis* and *P. porphyriacus* are the most cytolytic, while viper venoms showed weak membrane lysis as for *B. jararaca* to no membrane lysis as demonstrated by the venom of *C. horridus*.

## Discussion

The wealth of biological activities in venoms has attracted research in drug discovery and broader envenomation treatments. The complexity of venom mixtures urges for high content bioassays capable of identifying this diversity of bioactivities to improve studies exploring venoms. Advancements in high throughput cellular-based screens facilitated the exploration of venoms from spiders and scorpions which often lack cytolytic components and are rich in ion channels and receptors modulators^4,8,21^. Such approaches are not suitable for venoms rich in cytolytic components such as of snakes^1^, as cytolysis produces intracellular responses that can be confused with ion channel and receptors modulation^13^. This is a major caveat in venom-based drug discovery considering many venoms are a mixture of cytolytic and non-cytolytic bioactive entities, and therefore would benefit from high content bioassays to differentiate such activities early in the discovery process.

In this study, we developed a novel high content duplex assay combining calcium dye and propidium iodide that effectively detect and differentiate membrane lysis from ion pore formation and ion channels modulation responses simultaneously, and with a Z factor suitable to high-throughput (HTS) platforms. The combination of these two fluorescent dyes allowed detection of propidium iodide responses that is be attributed to its non-toxic and non-permeable characteristics ^22^, while the calcium 4 dye allowed specific detection of intracellular calcium attributed to its extracellular fluorescence masking technology (Molecular devices). Interestingly, the measurement of propidium iodide responses in the absence of calcium dye showed lesser fluorescence signals, indicating that the masking technology in the Calcium 4 dye is favouring the propidium iodide fluorescent signal. The simultaneous measurement of cytotoxicity and ion channel activity in HTS studies is a strong advantage in drug development as it speeds up the identification of leads with unwanted effects. In this study, we applied our duplex assay in combination with a Ca_V_ assay, and the voltage-gated ion channel activity was not affected by the presence of propidium iodide indicating the assay suitability to incorporation of screening for ion channels and receptors inhibitors such as voltage-gated sodium channels and acetylcholine receptors by adding a range of agonists for the desired targets under investigation^21,23^.

Evaluation of snake venoms revealed that *P. porphyriacus* and *P. australis* were cytolytic whilst *E. carinatus*, *B. jararaca* and *C. horridus* were not. The former was expected as *P. porphyriacus* and *P. australis* are Elapid species rich in 3FTXs with both cytotoxic and neurotoxic components ^24^. Elapid snakes are also rich in PLA2s which catalyse membrane phospholipids and causes membrane lysis ^25^. *E. carinatus* and *B. jararaca* belong to the Viperidae family, which contains PLA2s, SVMPs and SVSPs. The *E. carinatus* contains mostly PLA2s and SVMPs whereas *B. jararaca* contains an abundance in all of 3 proteins ^26, 27^. Given that PLA2s are highly cytotoxic, and that both these venoms showed low cytotoxicity in our bioassay, we concluded that the rapid membrane lysis and detection of DNA exposure is characteristic of Elapid venoms containing 3FTX cytotoxins and other non-enzymatic cytolytic components.

Intraspecific variation of *C. Horridus* existence of a neurotoxic variant (type A) and a heamotoxic variant (type B) ^28,29^. Type B is rich in SVMPs and SVSPs whereas type A is rich in SVSPs and PLA2s ^29^. It is possible that the crude venom sample from *C. horridus* was of type variant given no neuro and cytotoxicity was detected in our assay. The transient calcium response peak induced by the venoms of *B. jararaca* and *E. carinatus* could be associated to ion channel/receptors modulation or cellular stress while the propidium iodide response could be associated to a background response. The venom of the related viper *Bothrops moojeni* was found to induce intracellular calcium responses without cytotoxic effects in neuronal cells ^30^, which agrees with our results for *B. jararaca* and *E. carinatus*.

In conclusion, this is the first HTS high-content duplex assay of its kind. Further applications of these method can lead to novel discoveries in therapeutic and in the composition and evolution of animal venoms containing cytotoxic, cardiotoxic and neurotoxic components, as well as in the development of novel envenomation treatments.

## Data availability

All the raw data and derived data of this manuscript are available from the corresponding author upon reasonable request.

## Ethical Statement

No human subjects were involved in this work

## CRediT authorship contribution statement

**Fernanda Cardoso**: Conceptualization, Methodology, Supervision, Review and Editing, Funding **Simon Kramer**: Writing original draft, Investigation, Data curation, Validation, and Formal analysis **Charan Kotapati**: Investigation, Data curation, Validation, and Formal analysis **Yuanzhao Cao**: Investigation, Data curation and Formal analysis **Nathan Palpant, Bryan Fry and Glenn King:** Review and Editing, Resources.

## Acknowledgements

We acknowledge Dr Camila R Ferraz at the State University of Londrina, BR, for the venoms of the vipers B. jararaca and C. horridus.

